# An Automated 3D Analysis Framework for Optical Coherence Tomography Angiography

**DOI:** 10.1101/655175

**Authors:** Mona Sharifi Sarabi, Jin Kyu Gahm, Maziyar M. Khansari, Jiong Zhang, Amir H. Kashani, Yonggang Shi

**Affiliations:** USC Stevens Neuroimaging and Informatics Institute, Keck School of Medicine of University of Southern California, Los Angeles, CA, US; USC Roski Eye Institute, Department of Ophthalmology, Keck School of Medicine of University of Southern California, Los Angeles, CA, US

**Keywords:** Optical Coherence Tomography Angiography, 3D Vessel Skeleton Length, Diabetic Retinopathy

## Abstract

Optical Coherence Tomography Angiography (OCTA) is a novel, non-invasive imaging modality of retinal capillaries at micron resolution. While OCTA generates 3D image volumes, current analytic methods rely on 2D *en face* projection images for quantitative analysis. This obscures the 3D vascular geometry and prevents accurate characterization of retinal vessel networks. In this paper, we have developed an automated analysis framework that preserves the 3D geometry of OCTA data. This framework uses curvelet-based denoising, optimally oriented flux (OOF) vessel enhancement and projection artifact removal, as well as the generation of 3D vessel length from the Hamilton-Jacobi skeleton. We implement this method on a dataset of 338 OCTA scans from human subjects with diabetic retinopathy (DR) which is known to cause decrease in capillary density and compare them to healthy controls. Our results indicate that 3D vessel-skeleton-length (3D-VSL) captures differences in both superficial and deep capillary density that are not apparent in 2D vessel skeleton analyses. In statistical analysis, we show that the 3D small-vessel-skeleton-length (3D-SVSL), which is computed after the removal of the large vessels and associated projection artifacts, provides a novel metric to detect group differences between healthy controls and progressive stages of DR.

This work was supported in part by NIH grants UH3NS100614, R21EY027879, U01EY025864, K08EY027006, P41EB015922, P30EY029220, Research to Prevent Blindness, and UL1TR001855 and UL1TR000130 from the National Center for Advancing Translational Science (NCATS) of the U.S. National Institutes of Health.

## I. INTRODUCTION

Optical Coherence Tomography Angiography (OCTA) is a novel and non-invasive modality that is clinically approved to evaluate the retinal vasculature [1-3]. It has been applied in studying various ophthalmological diseases such as age-related macular degeneration (AMD), diabetic retinopathy (DR), uveitis, vein occlusions, and glaucoma [4-9]. In addition, OCTA has been shown to be a valuable imaging modality to identify changes in the microvascular network caused by neurovascular diseases [10-13]. While OCTA imaging devices provide 3D image volumes, analytical methods primarily use 2D *en face* images for quantitation [5, 9]. Qualitative analysis of 3D volume rendering has demonstrated that there are significant advantages to 3D analysis of OCTA data [14, 15] but quantitative 3D analysis has not yet been demonstrated. To better utilize the information in the 3D OCTA image volume, we have developed an analysis framework to reconstruct and quantify the 3D representation of retinal vasculature from OCTA images.

Various analysis methods have been proposed for 2D *en face* images derived from the 3D OCTA volume. [16] used the low-pass filters for denoising and an adaptive thresholding for vessel binary map generation. Multi-scale Hessian filters were also applied to OCTA data in animal studies [17, 18]. Eladawi et al [19] presented an automatic system using a joint Markov-Gibbs model, a Naïve Bayes (NB) classifier, and a 2D connectivity filter for segmentation of 2D superficial and deep retinal maps of normal and diabetic eyes. In an animal brain study, Li et al [20] proposed an angiogenesis tracking approach which combined top-hat enhancement [21], optimally oriented flux (OOF) [22], and graph-search based brain boundary detection algorithms. To the *en face* image of the OCTA volume at each depth, Yousefi et al [23] developed a hybrid method for vessel segmentation by combining multi-scale Hessian filters and an intensity-based model.

There are several challenges to developing 3D methods for accurate representation and analysis of OCTA data. These include high noise level, projection artifacts, residual motion artifacts, and vessel discontinuity [24]. The noise level significantly interferes with the interpretation and visualization of vessels, mainly the small capillaries. The projection of large superficial vessels is one of the main OCTA imaging artifacts that affects the appearance and geometry of retinal vessels in deeper layers [9, 25]. Vessel shape deformations such as neovascularization, microaneurysms, and pathologies like edema add more challenges to the 3D segmentation of vascular network in OCTA images.

In this work, we propose an automated analysis framework to systematically address these challenges and enable 3D measurements from OCTA image volumes [26]. An overview of our framework is shown in **Fig 1**. The inputs to our method include the original OCTA volume, and a retinal layer segmentation provided by the OCT-Explorer software [27-29]. As a first step, 3D curvelet transforms [30, 31] are used for OCTA image denoising with parameters adapted for disease severity (**Fig 1(c)**). In the second step, vessel enhancement is achieved based on optimally oriented flux (OOF) [22] (**Fig 1(d**)). Binary segmentation of all vessels is obtained by applying Otsu’s global thresholding on the OOF response map of vessels (**Fig 1(e**)). Using the multiscale property of OOF, a three-step algorithm is then developed to isolate the large superficial vessels together with their associated projection artifacts, thus providing a classification of large and small capillary vessels (**Fig 1(f**)). Finally, a 3D skeleton of the small vessels (**Fig 1(h**)) is calculated based on the Hamilton-Jacobi method [32], which provides the 3D vessel-skeleton-length (3D-VSL) for quantitative analysis.

**Fig 1.**
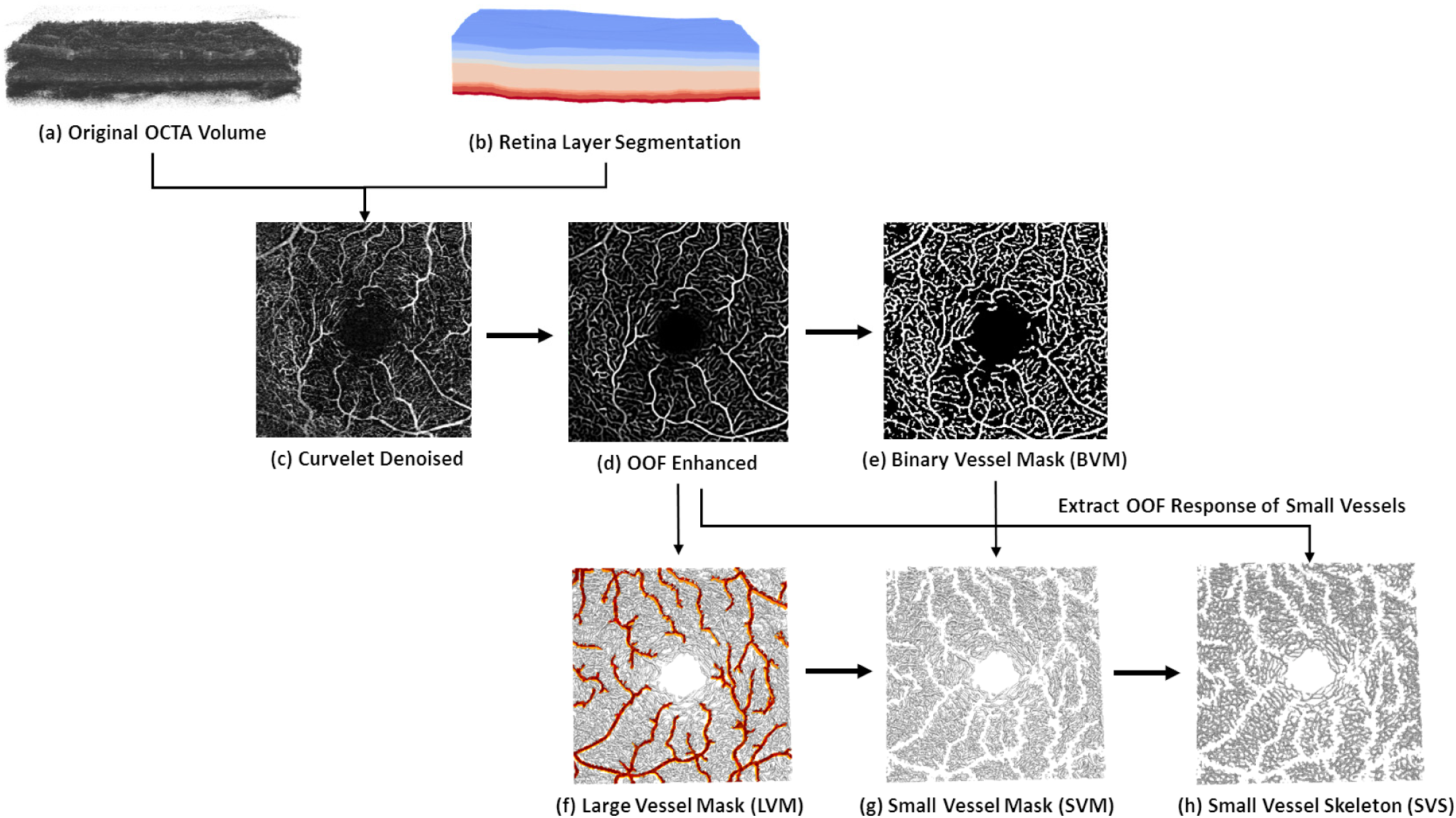
Method Overview. (a) Original OCTA image volume. (b) Retinal layer segmentation used to extract volumetric region of interest (ROI) which are layers 1 to 5 based on OCT-Explorer software. (c) Selected *en face* OCTA after applying 3D curvelet denoising on the ROI region of original OCTA image, OCTA ROI obtained by the inner product between(a) and (b). (d) Selected *en face* OCTA vessel map generated by computing the OOF response of (c). (e) Selected *en face* OCTA binary vessel mask (BVM) obtained by applying Otsu’s global thresholding on (d). (f) 3D large vessel mask (LVM) generated by applying a three-step OOF-based method on(d). (g) 3D small vessel mask (SVM) generated after removing (f) from (e). (h) Small vessel skeleton (SVS) obtained by the Hamilton-Jacobi method using (g) and OOF response of small vessels obtained by removing(f) from (d).

The rest of the paper is organized as follows. In Section II, we present the algorithmic details of the automated framework for 3D OCTA data. In Section III, we present experimental results from a dataset of 338 eyes from 230 subjects, consisting of healthy controls and subjects with various stages of diabetic retinopathy (DR). We demonstrate the feasibility of the proposed 3D method by quantifying 3D-VSL measures of superficial retinal layer (SRL), deep retinal layer (DRL), and retinal sublayers. In addition, we compare these results with 2D-based measures to illustrate the potential advantages of 3D-based analysis. Finally, discussions and conclusion are made in Section IV.

## II. METHODOLOGY

### A. Preprocessing

In this work, our main interest is to quantify the 3D features of retinal vessels which are significantly impacted in retinal vascular diseases such as diabetic retinopathy. Therefore, we first define a region of interest (ROI) in the OCTA scan limited to the neurosensory retina by excluding regions below the retinal pigment epithelium (RPE) and above the internal limiting membrane (ILM). This is achieved by defining the ROI based on structural layer segmentation of the OCT scan generated by the OCT-Explorer software [27-29]. Using a graph-based approach, the Iowa reference algorithm identified 10 surfaces in the retinal OCT scan as shown in **Fig 2**. The ROI for our analysis is defined as the volume from layer 1 to 5, i.e., the ILM to the outer plexiform layer (OPL). Furthermore, this ROI can be divided into superficial retinal layers (SRL) and deep retinal layers (DRL), which is composed of L1-L3 (ILM to inner plexiform layer (IPL)) and L4-L5 (Inner nuclear layer (INL)-OPL), respectively. According to the retinal anatomy, since L1 is typically very thin layer, L1 and L2 were merged in this study for the purposes of quantitative analysis and the results reported as the L2 sublayer in Section III of this paper represent the vessel measurements for L1+ L2.

**Fig 2.**
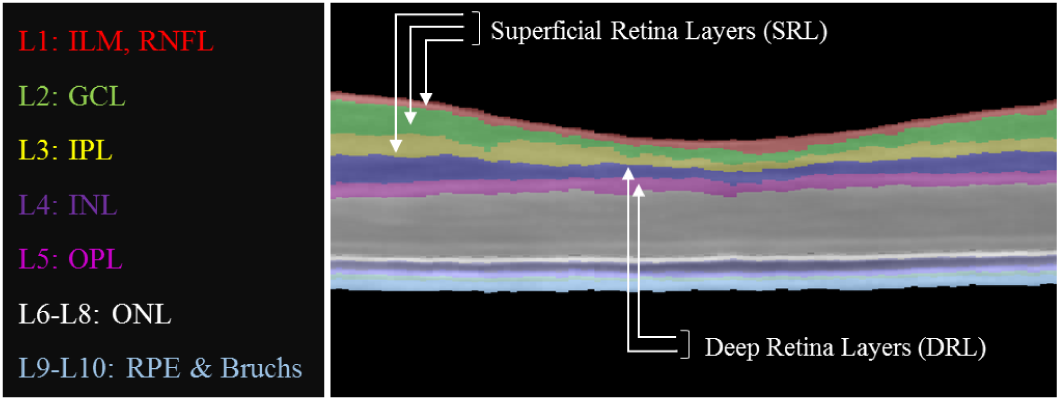
OCT structure layers. An illustration of the OCT layer segmentation achieved by OCT-Explorer.

To define the coordinate system in each OCT volume, we denote the direction along the A-scan as the z-axis, and the other two directions as the x- and y-axis. In the rest of this paper, x-y plane is also referred to as *en face* scan and x-z plane as the cross-section scan.

### B. Curvelet-based Denoising of 3D OCTA Images

The high noise level in OCTA images could cause the vessel discontinuity artifact that mainly affects the appearance of small vessels in OCTA scans. Conventional filtering methods generally obscure small vessels, limiting their feasibility for denoising 3D OCTA [5]. To match the denoising method with the curvilinear geometry of retinal vasculature in OCTA, we develop a curvelet-based OCTA denoising method [33].

Curvelet is a multiscale transform with high directional sensitivity and anisotropy characteristics due to its needle-shaped frame elements and provides an efficient representation of edges and other singularities along curves [30]. The three-dimensional curvelet transform has been proposed by Ying et al.[31]. In retinal image processing, the 3D curvelet transform were previously applied to OCT data to reduce the speckle noise [34]. Several properties of the curvelet transform make it an ideal candidate for OCTA denoising and vessel enhancement. The multiscale property of curvelets make them a proper choice for OCTA vessels, since they appear in different scales from the large superficial vessels to small capillaries in the deep retinal layers. Curvelets can be effectively used for elongated feature recovery, make it an appropriate tool for resolving OCTA vessel discontinuity. Anisotropy property of this transformation is also matched well with retinal vessel geometry. In this work, 3D Fast Discrete Curvelet Transform (FDCT) based on second-generation curvelet transform [35, 36] is used for OCTA denoising and vessel enhancement. The implementation is based on the wrapping method [30] proposed by CurveLab (http://www.curvelet.org). The detailed steps of the algorithm are shown in **Table I**.

**TABLE 1:**
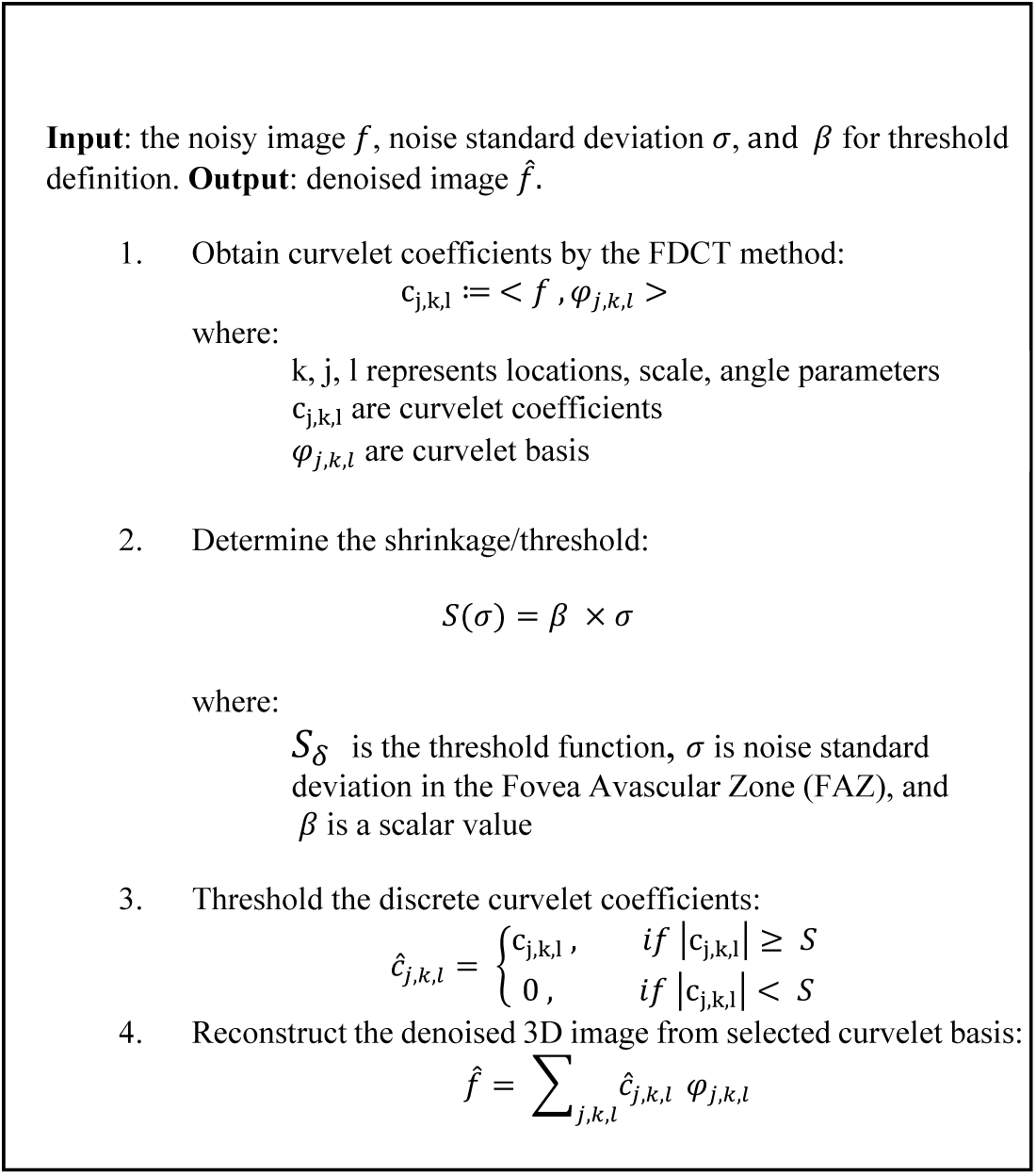
MAIN STEPS FOR CURVELET-BASED OCTA DENOISING

The most important parameters of curvelet-based denoising for OCTA are the number of scales and orientations and the threshold value for shrinking the curvelet coefficients. In all our experiments, the number of scales and orientations were empirically set to 6 and 16, respectively. The threshold value was defined as a function of noise standard deviation, which was estimated from the Fovea Avascular Zone (FAZ) of a set of representative OCTA images (**Fig 3(a), (b), (c), (d)**). Compared to the background area which is commonly used for noise estimation, the FAZ has similar texture and anatomical properties to neural retinal tissue with respect to light reflection and scattering, which is the basis of OCTA image construction. Therefore, the FAZ is a natural choice for noise estimation in inner retinal layers, or the defined ROI in this study. In order to define the FAZ, first the fovea center was selected (**Fig 3(b))**, as stated by[37], the avascular volume boundary of a normal eye is defined as 100 microns away from the fovea center in the x and y directions, expanded until the end of outer nuclear layer (ONL) in the z-direction (**Fig 2**). The 3D avascular mask was delineated using ITK-SNAP [38] on a set of ten representative OCTA images sampled equally from healthy controls and four DR stages. For pathological cases, we selected the subsets from the subjects that do not develop vessels in the foveal region and also have qualitatively preserved foveal morphology. The noise standard deviation was estimated to be 15 ± 1.5 from the FAZ intensity values of the selected OCTA scans. Since the observed values in selected scans were within the same range for both normal and DR scans, sigma was defined as 15 and the scaler value (β) was defined as 3 for all images in this study. This setting of parameters was validated on a variety of OCTA scans of healthy and DR patients. The results showed successful noise suppression in uniform areas while preserving small capillaries and resolving vessel discontinuity (**Fig 4(b)**).

**Fig 3.**
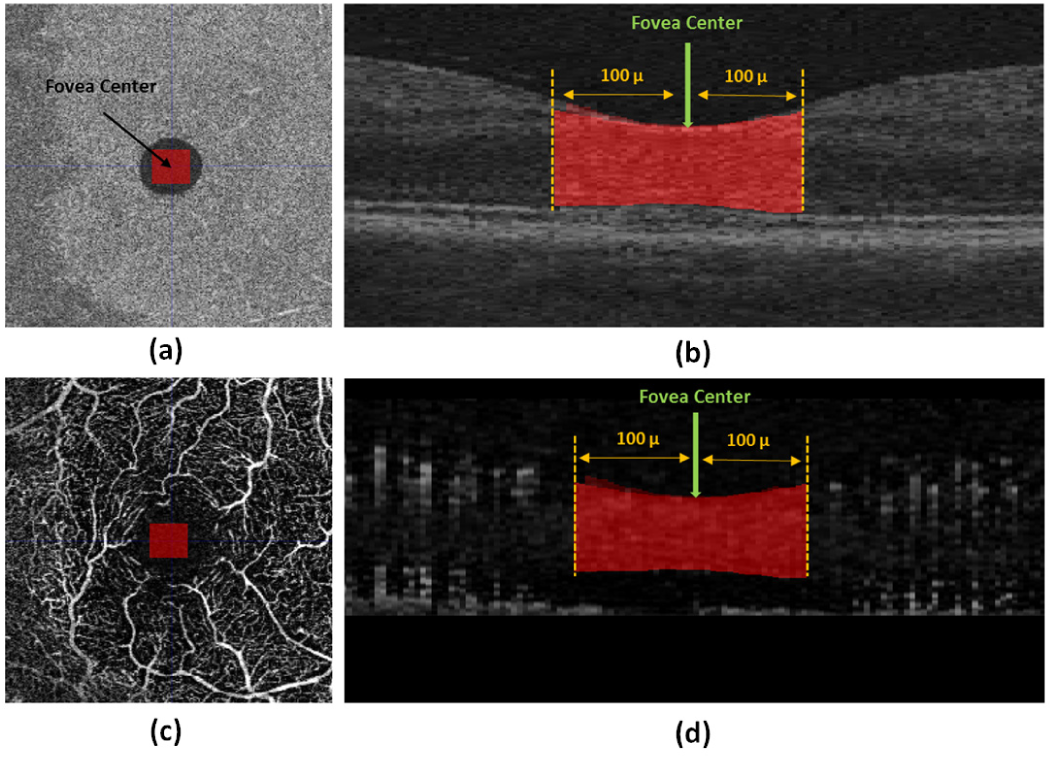
Estimation of noise standard deviation from the fovea avascular zone (FAZ). En face FAZ demonstration of (a) OCT scan, (c) OCTA scan of a healthy subject with 3mmx3mm FOV. Cross-section FAZ illustration of (b) OCT scan, (d) OCTA scan of the same subject.

**Fig 4.**
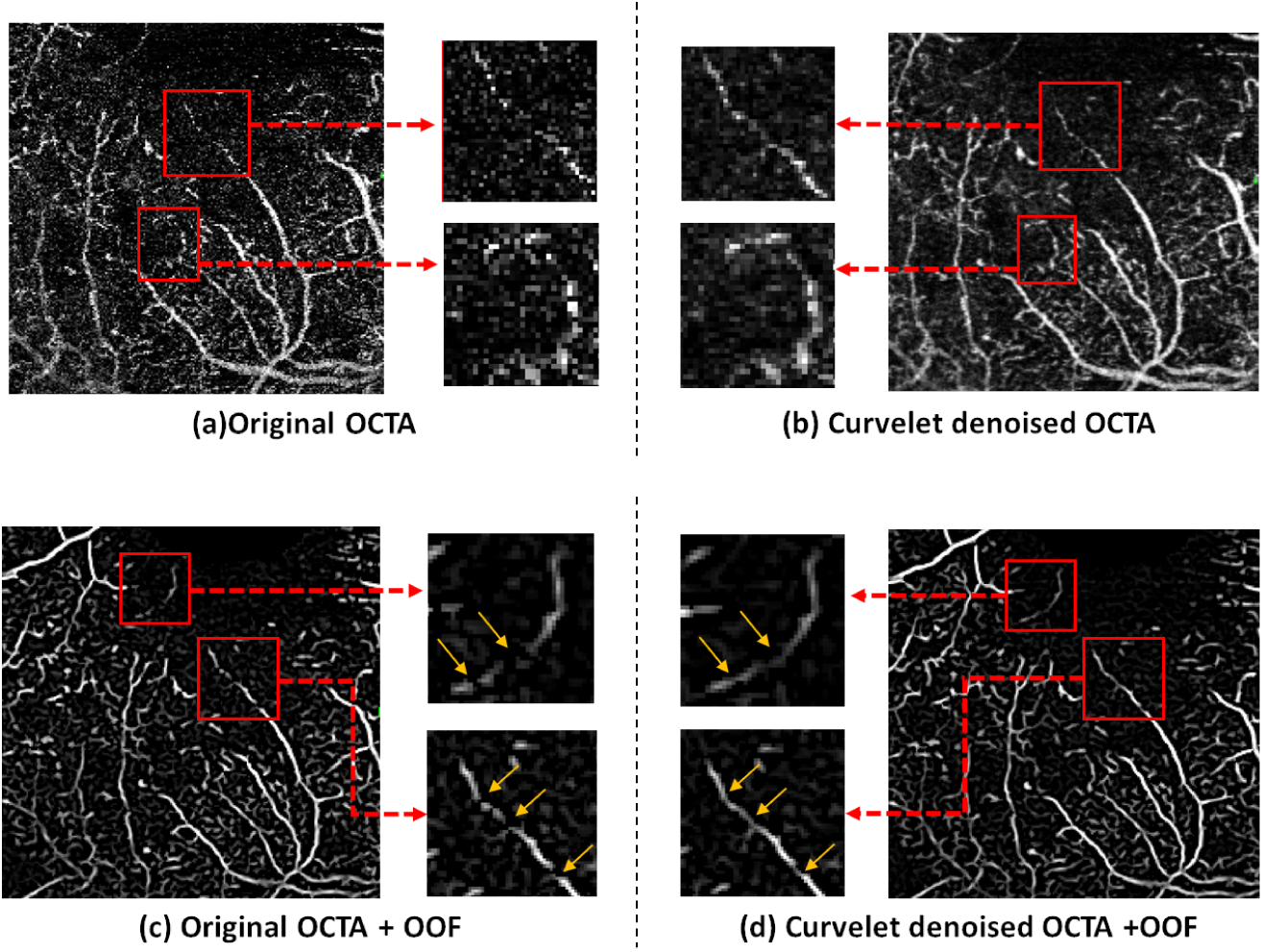
OCTA enhancement by Curvelet transform and OOF. Selected cross-sectional scan of a severe NPDR subject, (a) original OCTA. (b) Denoised OCTA using Curvelet transform. (c) OOF response obtained from original OCTA. (d) OOF response obtained from Curvelet denoised OCTA

### C. OOF-BASED VESSELNESS MAP

Vessel enhancement in 3D OCTA scans of retina is challenging due to the dense vessel network, closely located microvasculature, and inhomogeneous background. The Frangi Vesselness (FV) [39] and optimally orientated flux (OOF) [22] are among the most common filters used for vessel enhancement. Both filters provide multiscale curvilinear structure detection, but the OOF is capable of providing more accurate and stable detection responses [22] compared to the Hessian based method [40]. Therefore, we choose the OOF filter to provide an enhanced OCTA vesselness map because of its ability in detecting neighboring microvasculature structures in deep retinal layers while simultaneously suppressing the inter-vessel region.

The OOF response is generated for each voxel in the 3D image. At each image location X, the 3D OOF measure is calculated as the amount of the image gradient projected along a direction 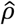 flowing out of a sphere *S*_r_ centered at the point X.

Formally, the OOF measure is define as an integral on the sphere:

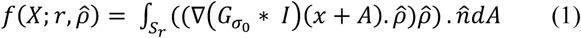

where 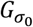 is the Gaussian kernel with σ_0_ as the standard deviation. The standard deviation is usually set to the voxel resolution to avoid interference from neighboring vessels. The variable *A* denotes the position vector on S_r_, dA as an infinitesimal area on S_r_, and 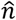 represents the outward unit normal of the sphere at *A*. The function 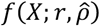 is the flux of the smoothed image gradient 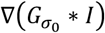 that is projected along 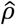 Using the divergence theorem, the quadratic form of *f* for a 3D image can be rewritten as:

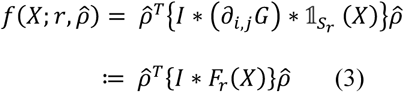

where ∂_i,j_*G* is the second derivative of G, 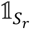 is the indicator function of the sphere S_r_ and F_r_ is the oriented flux filter. By differentiating the equation (3) with respect to 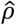, minimization of function f is in turn solved as a generalized eigenvalue problem. For 3D OCTA images, there are three sets of eigenvectors and eigenvalues. If the vessels are ideally represented as tubular shapes in the image, the first two eigenvalues would be much smaller than the third one, i.e., *λ*_1_(·) ≤ *λ*_2_(·) ≤ *λ*_3_ ≈ 0. Correspondingly, the first two eigenvectors span the vessel’s normal plane, and the third eigenvector represents the vessel orientation. Conventional OOF-based vesselness measures were defined with the product of the first two eigenvalues [22]. This is, however, not an appropriate choice for OCTA data. Fundamentally this is because the vessels do not necessarily appear tubular. Due to the presence of projection or tailing artifacts in the OCTA data, many vessels instead appear as plane-like structures (**Fig 5(a**)). In this case, the first eigenvalue will be much smaller than the second and third ones, i.e., *λ*_1_(·) ≤ *λ*_2_(·) ≈ *λ*_3_ ≈ 0. The first eigenvector will be along the normal direction of the plane-like structure, and the other two eigenvectors will span the plane. Based on these observations, we define the OOF-based vesselness measure for OCTA at each voxel as:

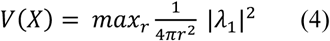

Since retinal vessels appear in different scales from large superficial layer vessels to tiny vasculature in deep layers, we used the multiscale property of the OOF vesselness measure in the above definition. To handle the varying scales of vessels, we follow the multi-scale approach of Law and Chung [22], which normalizes the eigenvalues of the OOF matrix by the surface area of the sphere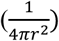. In this study, OOF-based vesselness enhancement is applied to the curvelet-denoised OCTA scans. The combination of these two techniques resolved both the vessel discontinuity and noise adjacent to tiny vessels while preserving the microvasculature geometry. **Fig 4 (c)** and **(d)** shows the comparison between OOF responses of an original OCTA image and an OCTA enhanced image by curvelet transform, respectively. The results show that OOF generates clearer and more continuous vesselness maps when applied after curvelet enhancement.

### D. LARGE VESSEL AND PROJECTION ARTIFACT REMOVAL

One of the main OCTA imaging artifacts is the projection artifact, also known as decorrelation tail (**Fig 5(a**)), that results in appearance of blood vessels in erroneous locations [25]. Multiple approaches were suggested for mitigating projection artifacts in OCTA images in 2D analysis based on *en face* images [37-39]. Using the OOF features above (**Fig 5(b**)), we developed a novel method for the removal of the large vessels from the superficial layer and their projection artifacts. This will allow us to focus on studying capillary changes in varying disease stages.

**Fig 5.**
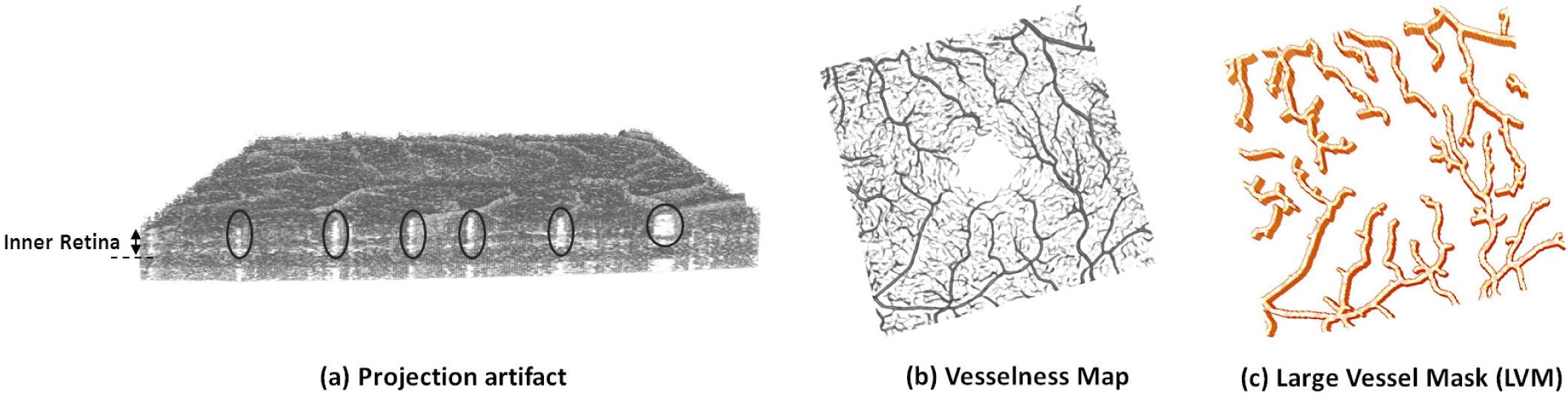
Projection artifact of large vessels and large vessels mask. (a) An illustration of projection artifact in the 3D OCTA image of a healthy eye. The black circles indicate the first type of projection artifacts caused by large vessels in superficial layers, which appear as elongated blood vessel shape of inner retina. (b) OOF vesselness map of the same subject (c) Large vessel mask (LVM) generated by our three-step algorithm based on OOF vessel features.

Using the multiscale property of OOF, we develop a three-step algorithm to construct the large vessel mask including their projection artifacts as illustrated in **Fig 5(c)**. Initially, large vessels were detected by two adaptive thresholding processes. Otsu’s global thresholding [41] was used to automatically choose the thresholds and applied first to the whole vesselness map to generate a binary vessel mask (BVM). After that, the same thresholding process was applied to the OOF-based vesselness map within the BVM to produce the large vessel mask (LVM). Then, we dilate the LVM along the z-direction of the OCTA image to include the projection artifact of large vessels from superficial layers. Finally, the LVM mask was dilated by one voxel along the x- and y-direction to ensure a complete inclusion of all voxels on the boundary of the large vessels, which might be missed from the initial thresholding process due to possibly low OOF response.

The resulting 3D BVM was postprocessed using connectivity analysis to remove small disconnected components. To find the major connected vessels, we determined the connected components included in the final vessel mask by measuring the volume of all components. Any outlier component with less than 1000 voxels was removed from the final vessel mask. Using the LVM and postprocessed BVM computed above, we can define a *small vessel mask* (SVM) as the set difference: BVM\LVM.

### E. VESSEL SKELETON

#### 1) 3D Vessels Skeleton

To compute the 3D Skeleton of OCTA vessels, we used the Hamilton-Jacobi skeletonization method [32]. The basic idea of this method is to extract skeleton points by their flux measure, followed by a homotopic thinning algorithm. Flux measure has large negative magnitude on the skeleton and is close to zero for the voxels away from the skeleton. In this paper, the flux measure was based on the negative OOF measure which had the same properties as the average outward flux measure of the vector field at each image point. To extract a thin and topology-preserving skeleton from the vessel masks, the flux-ordered and homotopic thinning process in [32] was applied. As can be seen from the examples in **Fig 7(d)**, the resulting skeletons follow the original vessel shape very well and provide the representation for vessel length calculation in 3D.

#### 2) En face Projection of Vessels Skeletons

In order to compare the accuracy of our method with conventional 2D analyses, we generated *en face* vessel skeleton images by calculating the maximum intensity projection of 3D vessel skeleton maps along the A-scan (z-direction). To enhance the comparability of these 2D *en face* images across subjects, OCT volumes were first brought to a common space by image registration. This was necessary since the orientation of OCT/OCTA volumes varies across scans. The registration was performed with the Elastix registration software [42] to compute a six parameter Euler transformation between a fixed volume and a moving volume. Note that we only correct for rotation and translation without stretching and scaling the OCT/OCTA images. For each OCT scan, a binary mask was first generated that includes retinal layers from Internal limiting membrane (ILM) to Bruch’s membrane (BM) by combining retinal layers segmented by OCT-Explorer. Zero padding (10 voxels along x and y-directions) was used to ensure no data is lost after the alignment. The binary mask of an ideal scan (well centered fovea and no image tilt) from a healthy subject was selected as the fixed volume and the atlas space for aligning other OCT scans. The binary masks from the rest of subjects were considered as moving volumes and registered to the fixed volume. Once the Euler transformation was computed, it was used to transform the 3D skeleton maps of each subject to the atlas space. Finally, the maximum intensity projection of the registered 3D skeleton map was calculated to construct the *en face* 2D skeleton map of each subject.

### F. EXPERIMENTAL RESULTS AND VALIDATIONS

#### 1) Dataset

All eyes included in this study were scanned at the USC Roski Eye Institute or affiliated clinics using a commercially available spectral-domain OCTA device (AngioPlex, Carl Zeiss Meditec, Dublin, CA, USA) with a scan speed of 68,000 A-scans per second, central wavelength of 840 nm and 3mm×3mm field of view. A fovea centered scan was taken in each eye of interest and generated an OCTA image volume of size 245×245×1024. Eye scans were included in the study if the OCTA signal strength was >7 and of sufficient quality for analysis. All images were manually reviewed for common imaging artifacts such as blockage of OCT signal by floaters and motion. Scans with floaters detected on the structural image were discarded if the decorrelation signal in the corresponding flow signal was also attenuated. Scans with more than 10 visible motion artifacts were also discarded. Diabetic retinopathy severity was graded based on clinical examination by board certified ophthalmologist with subspeciality training in retina. Scans from subjects with diabetes were categorized into mild nonproliferative diabetic retinopathy (NPDR), moderate NPDR, severe NPDR, and proliferative diabetic retinopathy (PDR) using the international classification of DR[43]. Patients with other retina vascular diseases were not included in this study. A total of 338 eyes remained in the study and their demographic data are shown in **Table II**.

**TABLE 2:** DATASET OVERVIEW

| Subject Characteristics | Healthy | Mild NPDR | Moderate NPDR | Severe NPDR | PDR |
| --- | --- | --- | --- | --- | --- |
| No. of subjects, n | 56 | 69 | 27 | 23 | 55 |
| No. of eyes (OD: OS) | 87(43: 44) | 109(60: 49) | 36(19:17) | 32(20:12) | 74(39:35) |
| Male:female | 43:44 | 70:39 | 18:18 | 14:18 | 38:36 |
| Age mean± SD (years) | 43±15 | 56±13 | 59±12 | 57±12 | 56±11 |
| Age range (years) | 21-75 | 23-80 | 31-91 | 32-79 | 29-83 |

### 2) Qualitative Assessment

In this experiment, we demonstrate our 3D OCTA analysis method on representative scans for the visualization and qualitative assessments of each intermediate step. **Fig 6** shows the results from the main steps of our method on an (x-y) plane of a severe NPDR(A1-A5) eye. In **Fig 6(a)**, A1 shows the original OCTA volume from the OCT machine, where the high noise level can be observed. A2 displays the same scan after curvelet-based denoising, and the result shows higher SNR and improved vessel connectivity as highlighted in **Fig 6(b)**, and **Fig 6(c)**. The panel A3 in **Fig 6(a)** is the result of OOF enhancement applied to the denoised scan from the previous step (A2). The outcome of this step demonstrates the effectiveness of OOF in preserving microvasculature while suppressing noise in closely located neighborhood. The binary vessel mask (BVM) shown in the A4 panel of **Fig 6(a)** was obtained by thresholding on the OOF response using Otsu’s method, where the small vessel mask (SVM) was generated by removing the large vessel mask (LVM) shown in red. Finally, the small vessel skeleton (**Fig 6(a)**: A5) was generated by the Hamilton-Jacobi method using the OOF response and the SVM. The 3D renderings of the analysis results from two representative examples (one normal and one severe NPDR) are shown in the first and second row of **Fig 7**, respectively. In each row, the original OCTA (**Fig 7(a)**), enhanced vessel network (**Fig 7(b**)), BVM (**Fig 7(c**)), and the small vessel skeleton (**Fig 7(d**)) are plotted. From these results, we can clearly see the morphologic differences between the vessel network of the normal and severe NPDR subjects. The normal subject has a dense vessel structure with a connected foveal ring, while the severe NPDR subject has sparse vessel network with vessel loss.

**Fig 6.**
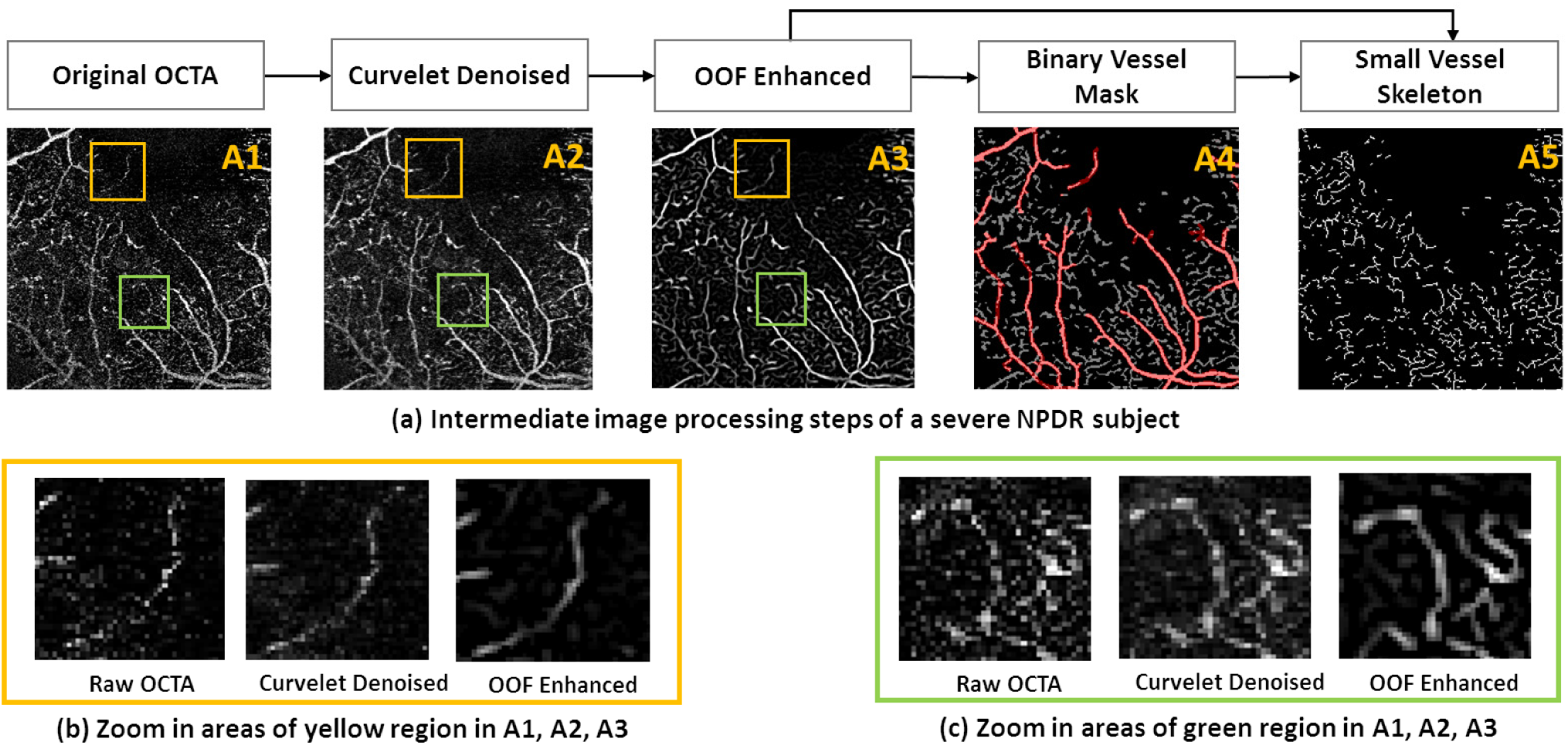
Main image processing steps of a selected *en face* plane of a severe NPDR eye. A1 illustrates *en face* scan of input OCTA from the OCT machine; A2 shows the same scan after reconstruction by curvelet transform; A3 represents the OOF response of A2; A4 is the vessel binary mask obtained by thresholding on the OOF response, large vessels mask shown in red removed using the proposed three-step algorithm; A5 is the skeleton of small vessels. (b), (c) Show the zoom-in areas of yellow and green boxes of A1, A2 and A3.

**Fig 7.**
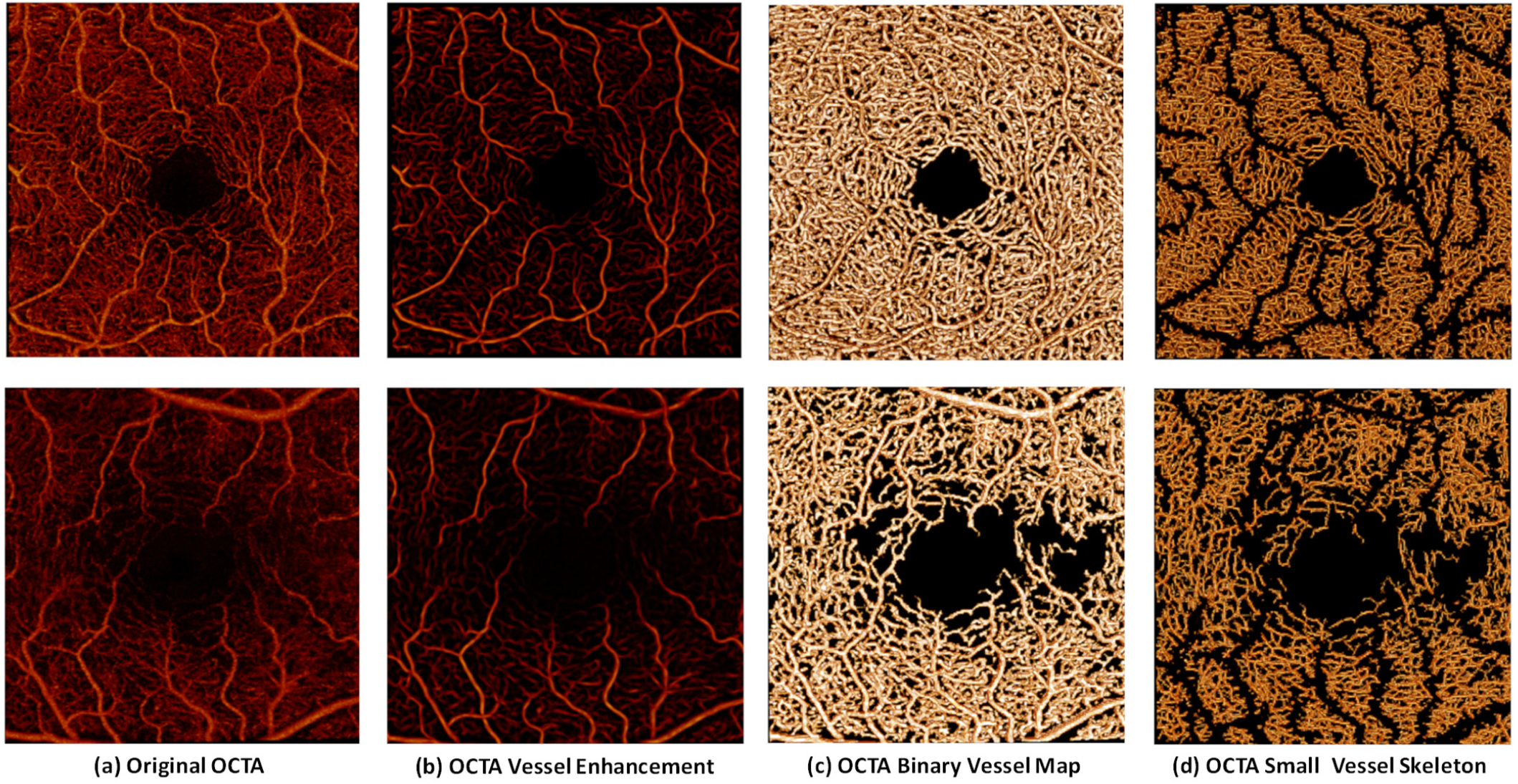
3D rendering of a normal (row1) and a severe NPDR (row2) eye. (a) Original OCTA. (b) Enhanced vessel map after curvelet-based denoising and OOF. (c) Binarized vessel mask. (d) Skeleton of small vessels obtained by the Hamilton-Jacobi method.

### 3) Quantitative Results

For quantitative analysis on the large-scale DR dataset, 3D vessel skeleton length was computed as the total number of voxels in the OCTA skeleton. The 3D vessel skeleton length was computed for all the vessels in the BVM, which we denote as all-vessel-skeleton-length (3D-AVSL), and the small vessels in the SVM, which we denote as small-vessel-skeleton-length (3D-SVSL). The 3D-AVSL and 3D-SVSL were calculated for the superficial and deep layers. As explained in section E-2 we also generated a 2D *en face* representation of these 3D images by counting the total number of pixels from an image obtained via maximum intensity projection of the 3D skeleton. This 2D *en face* representation of the 3D volumes was used to cross-validate our results with the conventional 2D *en face* measures described by our group and others in the past.

To demonstrate the potential benefits of 3D vs 2D vessel length measures, we conducted a quantitative comparison of vessel length measures from the two methods in the SRL and DRL using data of all healthy eyes. For both the 3D and 2D method, we computed the 3D-AVSL and 3D-SVSL and the results are shown in **Fig 8**. For both the 3D-AVSL and 3D-SVSL measures, t-test results indicate that SRL and DRL vessel length measures differ significantly based on our 3D analysis results, which are consistent with previous findings in histology [44]. On the other hand, the results from 2D projection of the skeleton image failed to show this difference.

**Fig 8.**
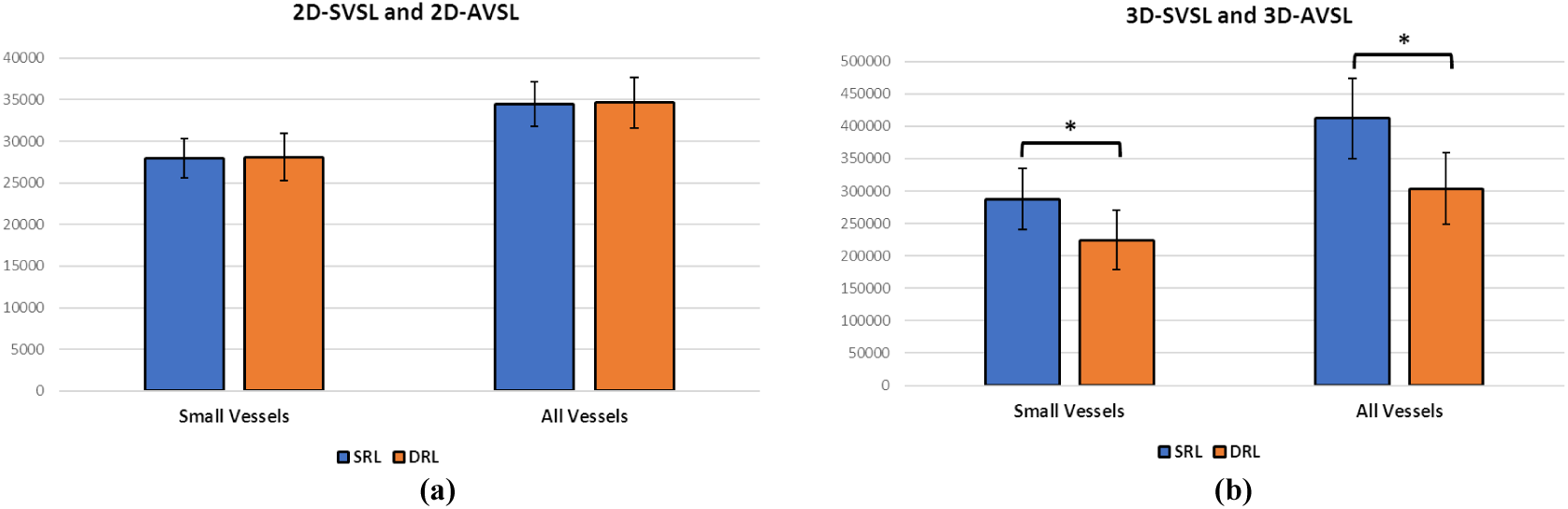
Comparison of quantitative OCTA results between 2D *en face* and 3D vessel measures of normal eyes. (a) bar charts illustrate the mean±SD of 2D small-vessel-skeleton-length (2D-SVSL) and 2D all-vessel-skeleton-length (2D-AVSL) in SRL and DRL of 2D *en face* image obtained via maximum intensity projection of the 3D skeleton. (b) bar charts illustrate the mean±SD of 3D-SVSL and 3D-AVSL.

Next, we applied our 3D method to study group differences between the controls and patients at different DR stages and between successive stages of DR. Statistical analysis was performed using SAS 9.4 (SAS Institute, Inc., Cary, NC). Multivariable linear regressions were used to compare the 3D-measures across groups. The generalized estimating equation (GEE) method was used to control for age, gender, and between-eye correlation when appropriate [6]. All statistical analysis was considered significant for p-value set to 0.05. As shown in **Fig 9(a)** and **Fig 10(a)**, both the 3D-AVSL and 3D-SVSL measures in SRL are significantly lower at all DR stages compared to healthy controls. In DRL, the pairwise comparison between the 3D-AVSL measures of healthy and DR eyes only had significant p-value for the most severe stage of DR (PDR). The 3D-SVSL successfully detects group differences between the healthy and DR eyes at earlier stages in the DRL.

**Fig 9.**
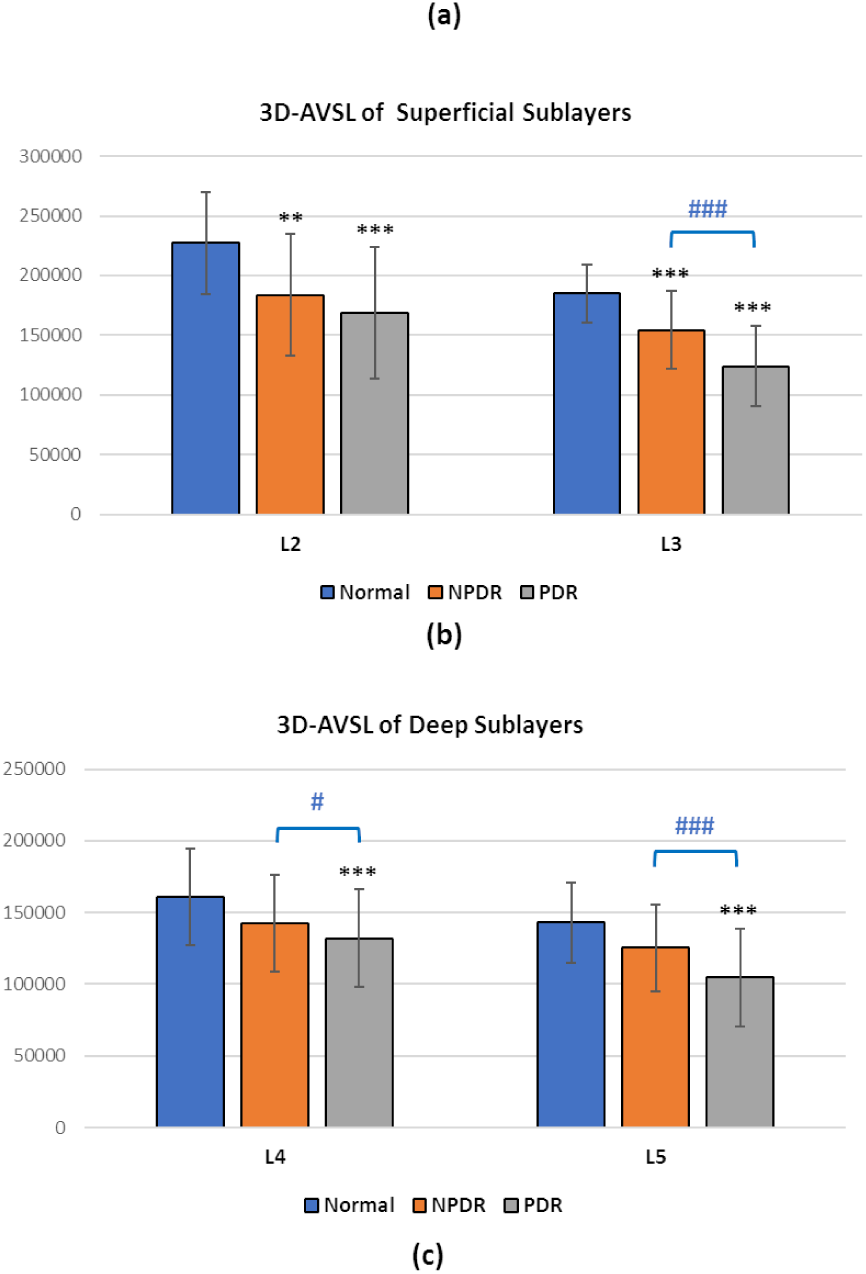
Comparison of OCTA quantitative results using 3D all-vessel-skeleton-length (3D-AVSL) measure between the controls and patients at different DR stages and between two successive stages of DR. (a) bar charts illustrate the mean±SD of 3D-AVSL in SRL and DRL. (b) bar charts illustrate the mean±SD of 3D-AVSL in superficial sublayers (L2: ILM-GCL; L3: IPL) of retina. (c) bar charts illustrate the mean±SD of 3D-AVSL in deep sublayers (L4: INL; L5: OPL) of retina. ‘*’ compares specified groups to controls. ‘*’, p<0.05; ‘**’, p<0.01; ‘***’, p<0.001.’ # ‘, compares specified DR group to the next stage of DR. ‘#’, p<0.05; ‘##’, p<0.01; ‘###’, p<0.001.

**Fig 10.**
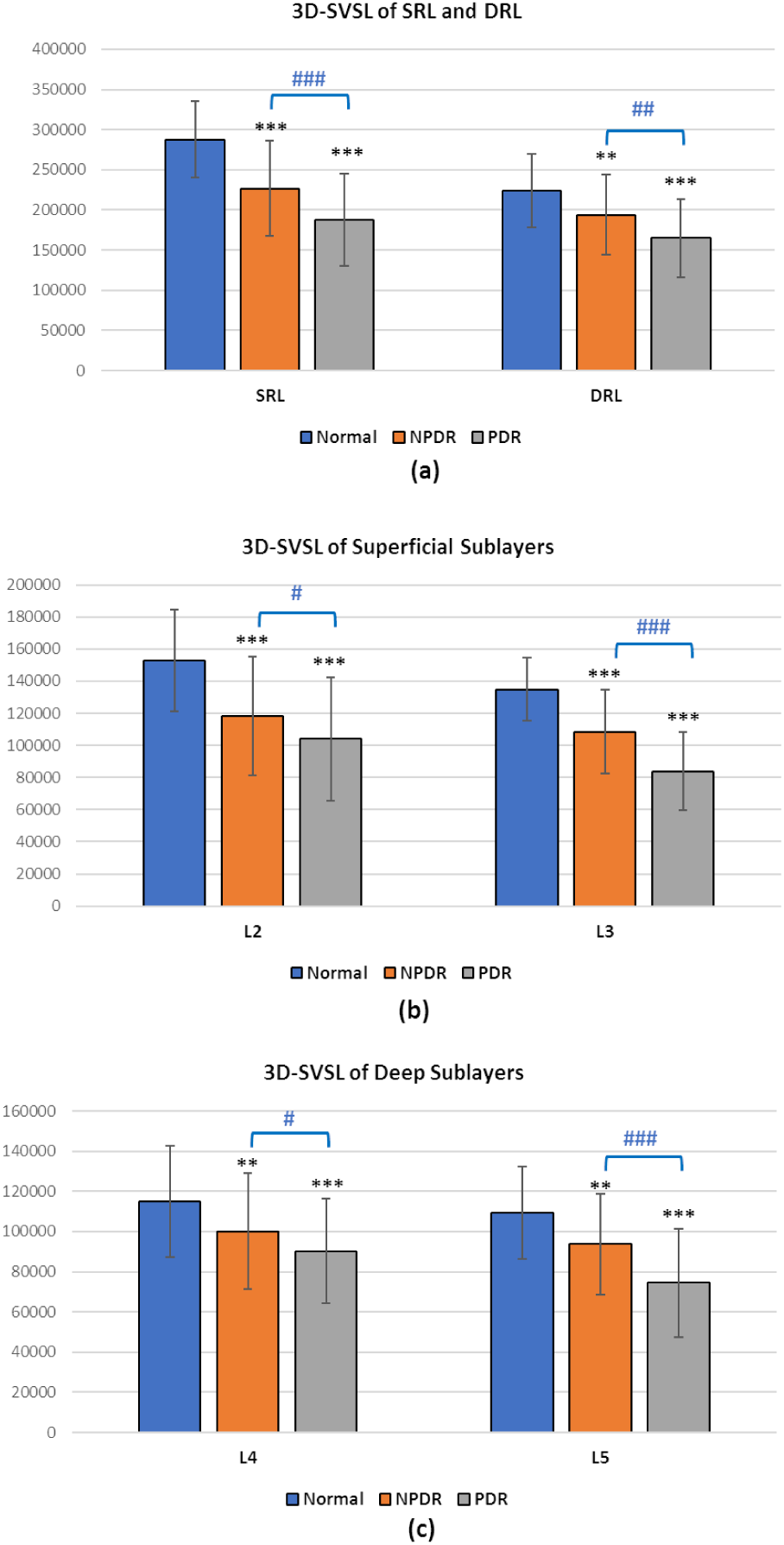
Comparison of OCTA quantitative results using 3D small-vessel-skeleton-length (3D-SVSL) measure between the controls and patients at different DR stages and between two successive stages of DR. (a) bar charts illustrate the mean±SD of 3D-SVSL in SRL and DRL. (b) bar charts illustrate the mean±SD of 3D-SVSL in superficial sublayers (L2: ILM-GCL; L3: IPL) of retina. (c) bar charts illustrate the mean±SD of 3D-SVSL in deep sublayers (L4: INL; L5: OPL) of retina. ‘*’ compares specified groups to controls. ‘*’, p<0.05; ‘**’, p<0.01; ‘***’, p<0.001.’ # ‘, compares specified DR group to the next stage of DR. ‘#’, p<0.05; ‘##’, p<0.01; ‘###’, p<0.001.

We also performed a sublayer analysis to evaluate the 3D measures efficacy in identifying more detailed group differences. While both the 3D-AVSL and 3D-SVSL achieved similar level of significance in detecting group differences in L3 (IPL) as shown in **Fig 9(b, c)** and **Fig 10(b, c)**, the 3D-SVSL is more effective in differentiating the group differences, especially between the healthy and the NPDR group, at the superficial L2 layer (ILM-GCL) and also deep L4 (INL) and L5(OPL) layers, which further demonstrates the value of removing the large vessels in the early detection DR-induced vasculature changes. Furthermore, the results of pairwise comparison between successive DR stages in the retina sublayers demonstrated that at the more superficial L2 layer, only 3D-SVSL was significantly different between NPDR and PDR subjects while 3D-AVSL did not capture this group differences.

### G. CONCLUSION

3D segmentation of retinal microvasculature in OCTA is difficult due to the high noise level and existence of imaging artifacts, mainly the projection of large superficial layers. Previous studies relied on the analysis of 2D *en face* projection of OCTA images. Although some studies have demonstrated the potential value of 2D metrics, they have the limitation of inevitably obscuring the morphological information in the original 3D vasculature networks of OCTA. In this work, an automated tool was developed for the 3D analysis of OCTA scans. As part of the method, we developed a novel OOF-based projection artifact removal algorithm to remove the large superficial vessels and their projection artifacts. In the proposed framework, 3D vessel length measures were developed to quantify retinal vasculature changes at different DR stage. Due to the flexibility of our method with respect to the selection of OCT structure layers, more detailed quantitative assessment can be performed for superficial, deep and sublayers of retina.

The proposed framework was validated using a large-scale dataset of healthy and DR eyes. Our results clearly demonstrate the value of 3D-based analysis and the removal of large vessels and their projection artifacts. Statistical results based on 3D-SVSL measures showed that both SRL and DRL metrics can distinguish healthy eyes from DR eyes at different stages of the disease, but the SRL measures are more effective in detecting these group differences. Furthermore, sublayer analysis demonstrates that the 3D-SVSL measure can more effectively differentiating the NPDR groups from the healthy controls and PDR.

For future work, we will extend our method to OCTA scans of different FOVs and data from different OCT scanners. We will also apply our method to examine the longitudinal vascular changes in population studies or tracking changes during treatment of edema pathology.

